# Seasonal variations of ^137^Cs concentration in freshwater charr through uptake and metabolism in 1–2 years after the Fukushima accident

**DOI:** 10.1101/2021.05.28.446093

**Authors:** Kengo Okada, Masaru Sakai, Takashi Gomi, Aimu Iwamoto, Junjiro N. Negishi, Masanori Nunokawa

**Affiliations:** Graduate School of Agriculture, Tokyo University of Agriculture and Technology, Tokyo, Japan; Fukushima Regional Collaborative Research Center, National Institute for Environmental Studies, Japan, Fukushima, 963-7700, Japan; Faculty of Environmental Earth Science, Hokkaido University, Hokkaido, Japan; Civil Engineering Research Institute for Cold Region, Hokkaido, Japan

**Keywords:** freshwater fish, leaching, radiocesium, resource subsidy, riparian forest

## Abstract

Understanding the factors influencing ^137^Cs concentrations in freshwater salmonids is crucial for reviving inland fisheries in polluted regions. We studied seasonal variations of ^137^Cs concentration in charr (*Salvelinus leucomaenis*) through uptake and metabolism in forested headwaters at Fukushima and Gunma sites. Charr consumed both terrestrial and aquatic animals, and terrestrial prey was predated more in summer at both sites. The ^137^Cs concentrations in litter, which is a dominant basal food resource of both forest and stream ecosystems, differed between forest and stream due to ^137^Cs leaching effect on litter submerged in streams. The concentration difference in both litter and prey was greater at Fukushima site than at Gunma site. The estimated prey ^137^Cs concentration at Fukushima site peaked in summer when terrestrial preys are most available, whereas it remained relatively constant at Gunma site because of the small difference of ^137^Cs concentrations in between terrestrial and aquatic preys. The specific metabolic rate of charr was commonly changed with stream water temperature, greatest in summer and lowest in winter at both sites. Because both prey ^137^Cs concentrations and specific metabolic rates peaked in summer, the combination of uptake and metabolism at Fukushima site largely negated seasonal ^137^Cs fluctuations in charr, whereas specific metabolic rate fluctuations could be the major determinant of charr ^137^Cs concentrations at Gunma site. Our results suggested that ^137^Cs concentrations in prey items, whose seasonality are varied due to initial ^137^Cs fallout volume, were expected to be an important determinant for ^137^Cs concentrations in charr.

## 1 INTRODUCTION

Headwater streams harbor freshwater salmonids including *Salvelinus* and *Oncorhynchus* species in temperate montane forests of Japan (Masuda et al. 1984). Although these species are popular for local freshwater fisheries, the Fukushima nuclear accident occurred in 2011 has caused serious ^137^Cs accumulations in the fishes around Fukushima due to the heavy burden of bioavailable ^137^Cs provided by surrounding forests (Sakai et al., 2021; Wada et al., 2019).

Freshwater fishes take up cations, including ^137^Cs, largely through feeding and have lower cation excretion rates during osmoregulation than marine fishes (Wada et al., 2019). Thus, the composition of food items and their ^137^Cs concentrations are important determinants of ^137^Cs concentrations in headwater salmonids (Haque et al., 2017; Matsuda et al., 2020). In addition, because ^137^Cs elimination through metabolism also influences ^137^Cs concentration in fish (Doi et al., 2012; Hessen et al., 2002; Holloman et al., 1997; Kolehmainen, 1972; Peles et al., 2000; Rowan et al., 1997), dynamics of ^137^Cs concentrations in headwater salmonids needs to be examined by considering both processes involved in uptake and metabolism.

Headwater salmonids consume both terrestrial and aquatic prey, depending on seasonal availability (Nakano et al., 1999a; Nakano & Murakami, 2001). In riparian forest and stream ecosystems, because detrital food webs originating from plant litter are of most potent energetic pathways (Gomi et al., 2002; Miyashita et al. 2003; Sakai et al., 2016a; Vannote et al., 1980; Wallace et al., 1997), the detrital food webs may promote ^137^Cs transfers from contaminated litter to salmonids through potential detritivorous preys (Negishi et al., 2018). Meanwhile, previous studies found that ^137^Cs in litter greatly leach after falling into streams (Gomi et al., 2018; Sakai et al., 2015; 2016b). The ^137^Cs leaching effect causes greater relative difference of ^137^Cs concentration in litter between terrestrial and aquatic environments in more contaminated sites (Sakai et al., 2016b), and the concentration difference results higher ^137^Cs concentrations in forest animals than in stream ones in heavily contaminated sites (Sakai et al., 2016a). The suite of the studies implies that terrestrial prey for salmonids may be more contaminated compared to aquatic prey in headwaters which suffered heavier initial ^137^Cs fallout.

Because metabolisms of fish are largely controlled by temperature (Brown, 2004), ^137^Cs elimination through metabolism will be higher in summer and lower in winter. Meanwhile, terrestrial prey for salmonids are more available in summer than in winter (Baxter et al., 2005; Nakano et al., 1999a). Therefore, overall ^137^Cs concentration in preys may peak in summer in heavily contaminated sites where ^137^Cs leaching from litter induces apparent difference of ^137^Cs concentrations in between terrestrial and aquatic preys. In contrast, overall ^137^Cs concentration in preys may not show clear seasonality in less contaminated sites. These hypotheses suggest that ^137^Cs leaching effect on aquatic litter, which varies with degree of initial ^137^Cs fallout, influences seasonality of ^137^Cs uptake and consequently ^137^Cs concentrations in salmonids.

In this study, we examined how different initial fallout volumes influence seasonal changes in ^137^Cs concentrations in freshwater charr (*Salvelinus leucomaenis*) based on temporal dynamics in uptake and metabolism. The result highlights the importance of ecological processes driving contaminants’ transfer across ecosystems through resource subsidy (cf. Walter et al., 2008). This would provide useful information for the contamination management of freshwater salmonids in headwater streams that suffered ^137^Cs fallout.

## 2 MATERIALS AND METHODS

### 2.1 Study site

This study was conducted in headwater streams and surrounding riparian zones in Fukushima and Gunma Prefectures, Japan. The Fukushima and Gunma sites belong to the Abukuma River and Tone River systems, respectively. The Fukushima site was located approximately 45 km west of the Fukushima Daiichi nuclear power plants (FDNPPs) (37°36’N, 140°37’E), and the Gunma site was located 180 km southwest of the FDNPPs (36°33’N, 139°21’E). Based on airborne observations conducted in June 2012, ^137^Cs deposited on the ground surface was 100–300 kBq/m^2^ at the Fukushima site and 30–60 kBq/m^2^ at the Gunma site (MEXT, 2012). The Fukushima and Gunma sites have drainage basins of 170 and 93 ha, respectively. Based on AMeDAS automated weather station data collected nearby, the Fukushima site had mean annual precipitation and air temperature of 1248.4 mm and 10.9°C. Based on another nearby weather station, the Gunma site had mean annual precipitation and air temperature of 1323.3 mm and 14.5°C. The channel reaches had typical step-pool morphologies (Montgomery & Buffington, 1997). The riparian zones of both study sites were covered with evergreen conifer (*Cryptomeria japonica*) plantation that is a dominant forest type of Japan. We selected 50-m study areas along the stream channels and 20-m-wide areas in the riparian zones on both sides of the channels. The wetted widths of the study areas at both sites were similar (1–5 m).

### 2.2 Field investigations

We sampled charrs, preys and litter four times, in summer (23–25 August for Fukushima and 16–18 August for Gunma) and autumn (20–22 November for Fukushima and 13–15 November for Gunma) in 2012, and winter (19–21 February for Fukushima and 14–16 February for Gunma) and spring (16–18 May for Fukushima and 26–28 May) in 2013. Charrs were collected using a backpack electrofisher (LR-24; Smith-Root, Inc., Vancouver, WA, USA), measured their wet weight and standard body length, and collected stomach content samples from each fish with stomach pumps. Charr was the only fish species inhabiting in both study streams, with densities at the Fukushima and Gunma sites of 0.19 and 0.28 individuals/m^2^, respectively, estimated based on the three-pass removal method. The samples were collected in the early morning and at dusk, when prey was potentially most available (Nakano et al., 1999b). The stomach content samples were immediately preserved in 99.5% ethanol. To avoid local extinctions of the charr populations, we only sacrificed randomly selected 3–8 individuals at both sites and from all seasons for measurements of ^137^Cs concentrations in charr (20.5% and 11.5% of total catch at the Fukushima and Gunma sites, respectively, Figure 1). The charr samples were stored at -20°C prior to laboratory analyses.

**FIGURE 1.**
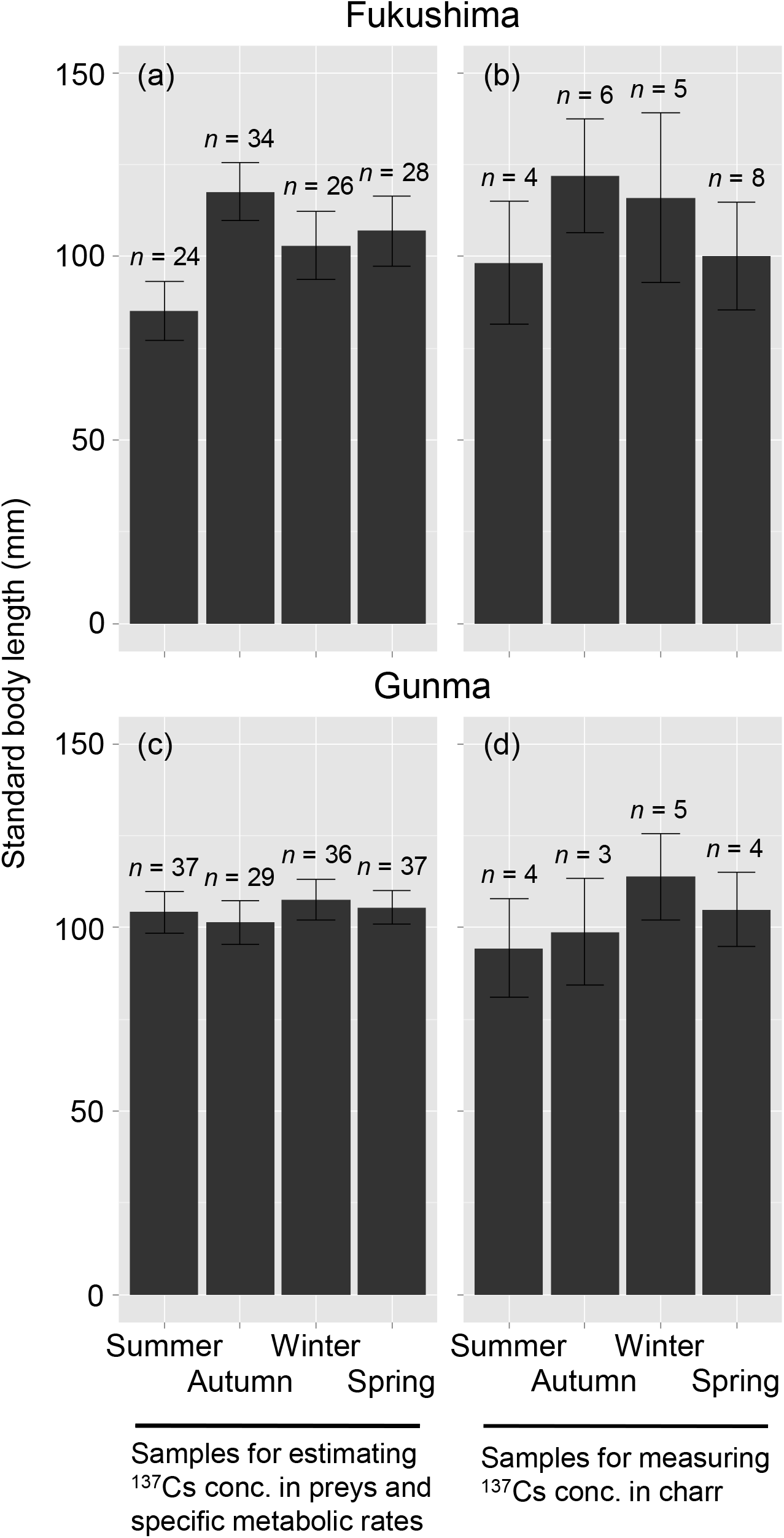
Standard body lengths of charr samples of this study. The standard body lengths of the charr samples used for estimating ^137^Cs concentrations in prey and specific metabolic rates at the Fukushima and Gunma sites (a and c). Those used for measuring ^137^Cs concentrations in charr of the Fukushima and Gunma sites (b and d). The sample sizes are shown at tops of each bar. Error bars indicate ± standard error. Summer: August, autumn: November, winter: February, spring: May.

To determine potential aquatic and terrestrial prey, we sampled stream channels with driftnets (25 × 25 cm; mesh size, 250 μm) and riparian zones with pan traps (1.05 × 0.60 × 0.21 m). Five driftnets were placed longitudinally in each stream at 10-m intervals for 1 h in the early morning and at dusk in each season. The volume of water flowing through the driftnets was estimated from the water depth and current velocity measured using an electromagnetic current meter (VE20; Kenek Co., Tokyo, Japan) at the openings of the driftnets. Five pan traps filled with several centimeters of water and a few drops of surfactant were placed along the study reaches at 10-m intervals for 7 days. During the sampling periods in autumn and winter, antifreeze liquid was added to the pan traps to prevent the water from freezing. The samples obtained from the driftnets and pan traps were preserved in 99.5% ethanol immediately after collection.

Measuring ^137^Cs concentrations in individual invertebrates such as insects is typically difficult due to their small individual body mass. Therefore, it was necessary to collect many individuals and pool samples by taxon to provide sufficient sample volumes for the radionuclide measurements (≳ several grams in dry weight) besides the above-mentioned investigations using stomach pumps, driftnets and pan traps. Benthic macroinvertebrates representing aquatic prey were collected from the study sites using D-frame nets. The collected macroinvertebrates were handled with tweezers and stored in glass vials grouped by taxon visually identified to the lowest possible taxonomic rank. The all collections were done after fish investigations to avoid anthropogenic disturbance which may affect foraging of charr. Invertebrates inhabiting the forest floor representing terrestrial prey were collected with 14 baited traps (diameter, 8 cm; depth, 11 cm) located 5–10 m from the sides of the streams. We filled the traps with an attractant consisting of a 1:1 admixture of beer and lactic acid beverage (Sakai et al., 2016a) and left them on the forest floor for 36 h. The collected samples were gently washed with distilled water, visually identified to the lowest possible taxonomic rank, and stored in plastic bags. In addition, spiders, frogs, and crabs were sampled by hand within the study riparian zones during each sampling period. All the organism samples were stored at -20°C prior to laboratory analyses.

To estimate the ^137^Cs concentrations of the basal food resource of the streams and riparian zones, we collected Japanese cedar litter deposited on the forest floors and in the streams of the study areas. Previous carbon and nitrogen stable isotope analyses had identified that terrestrial and aquatic food webs in Japanese cedar plantations originate mainly from Japanese cedar litter because of its persistence and abundance (Sakai et al., 2016a; 2016c); therefore, we assumed that the basal food resource of our study ecosystems was Japanese cedar litter. To avoid variations in ^137^Cs concentration due to decomposition stage, we collected intact litter both in the riparian forests and streams. Three litter samples were collected for both terrestrial and aquatic environments in each season at the sites. Stream water temperature was monitored in both stream channels every 10 min using temperature loggers (TidbiT V2; Onset Computer Co., Bourne, MA, USA) to calculate specific metabolic rates of charr (see 2.5).

### 2.3 Laboratory analysis

Muscle tissues of the charr samples were extracted and dried at 60°C for 2 days. The terrestrial and aquatic prey samples for radioactivity measurements were dried at 60°C for more than 2 days. Japanese cedar litter was dried in a similar manner. All dried samples were ground into powder using either an agate mortar and pestle or electrical mill (Osaka Chemical Co., Ltd., FM-1). The samples were packed into 100-mL polystyrene containers and sealed until the radioactivity measurements.

Prey sampled from stomach pump, pan traps, and driftnets were identified to the order (excepting terrestrial Oligochaeta and Diplopoda) and separated into terrestrial or aquatic organisms. Then, we considered adult aquatic insects to be aquatic animals due to the dominance of aquatic environments in their life history (Merritt et al., 2008), and frog species were included as terrestrial animals because they mainly predate terrestrial invertebrates (Maeda & Matsui, 1999). The samples were dried at 60°C for more than 2 days, and the dry weights were measured within 0.01 mg. From this, we estimated the total, terrestrial, and aquatic prey biomass of charr. Prey availability was expressed as biomass per volume of water flow (mg/m^3^) for the driftnet samples, biomass per area per day (mg/m^2^/day) for the pan trap samples, and biomass per individual (mg/individual) for the stomach pump samples.

### 2.4 Radioactivity measurements

^137^Cs radioactivity was analyzed in charr, prey, and litter using gamma ray spectroscopy. Gamma ray emissions at 661.6keV were measured using a high-purity germanium coaxial detector system (GEM20-70; Ortec, Oak Ridge, TN, USA) coupled with a multi-channel analyzer (DSPECjr2.0; Ortec). The energy and efficiency calibrations were performed using standard and blank (background) samples. The geometry was held constant when counting all samples for ^137^Cs concentrations. For the ^137^Cs activity analysis, all samples had counting errors <10%. All activities were corrected for decay based on the sampling date, and ^137^Cs concentrations below the detection limit were designated as 0 Bq/kg for the statistical analysis. The detector system was assessed using proficiency test samples from the International Atomic Energy Agency (IAEA-TEL-2012-03).

To estimate representative values for ^137^Cs concentrations in terrestrial and aquatic preys, we calculated the median ^137^Cs concentrations for them collected in all the sampling periods. Then, sum of multiplication of the median concentrations and proportions of biomass of terrestrial and aquatic prey in the stomach contents was used as “estimated ^137^Cs concentrations in prey” for individual charr. Because ^137^Cs concentrations in animals can differ between terrestrial and aquatic environments due to ^137^Cs leaching from litter (Sakai et al., 2016a), the relative proportions of terrestrial and aquatic prey could substantially influence ^137^Cs uptake in charr.

### 2.5 Specific metabolic rate

Specific metabolic rate is an important parameter for estimating whole-body ^137^Cs concentrations in fish (Ugedal et al., 1992) that was used in this study as an indicator of ^137^Cs elimination from charr. The specific metabolic rate (*B*) (W/kg) of charr was calculated based on the equation below (Brown 2004):

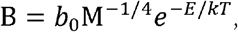

where *b*_0_ is a normalization constant independent of body size and temperature, *M* is the wet weight of charr (kg), *E* is the activation energy, *k* is Boltzmann’s constant, and *T* is the absolute water temperature (K). The wet weights of each charr were substituted into *M*, and *T* was obtained from the average of the water temperatures recorded over 1 week at the start of the field investigation.

### 2.6 Statistical analysis

One-way analyses of variance (ANOVAs) with permutation test (Legendre, 2007) were used for testing differences among seasons for (1) standard body lengths of charr in both samples for estimating ^137^Cs concentrations in preys and specific metabolic rates and for measuring ^137^Cs concentrations in charr, (2) prey biomasses from terrestrial and aquatic environments estimated with driftnets, pan traps, and stomach pumps, (3) estimated ^137^Cs concentrations in prey, (4) specific metabolic rates of charr, and (5) ^137^Cs concentrations in charr for each site. When significant, we used the same permutation ANOVA test for post hoc comparisons among the pairs of seasons. The permutation test allowed us to correctly handle the unbalanced data set of the study. We adjusted p values for multiple tests using the false discovery rate correction to avoid inflated type I error rates.

One-way ANOVAs with permutation test was also used to test differences of standard body length of charr between the samples for estimating ^137^Cs concentrations in preys and specific metabolic rates and for measuring ^137^Cs concentrations in charr. Further, we tested the difference in ^137^Cs concentrations in litter and prey between habitats (i.e., terrestrial and aquatic) using the ANOVAs. All statistical procedures were conducted using R 3.6.3 (R Core Team, 2020) with anova.1way function (Legendre, 2007) and rcompanion package (Mangiafico, 2020).

## 3 RESULTS

### 3.1 Body size distribution of charr

The comparisons among seasons indicated that standard body lengths of charr were similar in both the samples for estimating specific metabolic rate and ^137^Cs concentrations in prey and for measuring ^137^Cs concentrations in charr (Table 1 and Figure 1a–d). Also, the size distribution was similar between the two sample categories. Although positive correlations between ^137^Cs concentration and body size were reported in fishes (Ishii et al., 2020; Wada et al., 2019), we assumed that such body size effect is negligible for further discussion in this study.

**TABLE 1.** Results of permutation analyses of variance testing for effect of season on the standard body lengths of charr (sample 1: the samples for estimating specific metabolic rate and ^137^Cs concentrations in preys, sample 2: the samples for measuring ^137^Cs concentrations in charr) and comparing standard body lengths of charr between the samples 1 and 2.

|  | Fukushima |  | Gunma |  |
| --- | --- | --- | --- | --- |
| Factor | *F* | *P* | *F* | *P* |
| Sample 1: |  |  |  |  |
| Season | 2.27 | 0.08 | 0.21 | 0.90 |
| Sample 2: |  |  |  |  |
| Season | 0.38 | 0.78 | 0.69 | 0.54 |
| Sample (1 vs 2) | 0.03 | 0.86 | 1.07 | 0.33 |

### 3.1 Charr prey

The driftnet and pan trap samples indicated that prey tended to be more abundant in summer and spring than in the other seasons at both the Fukushima and Gunma sites (Table 2). Seasonal variations in terrestrial and aquatic prey biomass were similar and peaked in summer and spring, though the statistical significances were detected only in some pan trap data (Tables 2 and 3). Based on the stomach content samples, charr consumed terrestrial prey more in summer and aquatic prey more in autumn and spring (Table 2). Charr at both the Fukushima and Gunma sites consistently consumed greater amounts of prey in summer, autumn, and spring and less in winter, though there was no statistically significant difference among seasons (Table 2). The three most common terrestrial taxa found in the stomach content samples from fish at the Fukushima site were Anurans (36.3%), Orthopterans (22.6%), and Oligochaetans (15.3%), whereas those at the Gunma site were Orthopterans (40.2%), Coleopterans (25.7%), and Anurans (16.3%) (Table 4). The Orthopterans found in the guts were considered as taxa of Rhaphidophoridae in both the sites. The three main aquatic taxa in the stomachs of the fish at the Fukushima site were Isopods (33.9%), Decapods (28.3%), and Plecopterans (10.4%), whereas those at the Gunma site were Trichopterans (26.7%), Dipterans (24.8%), and Decapods (14.7%) (Table 4).

**TABLE 2.**
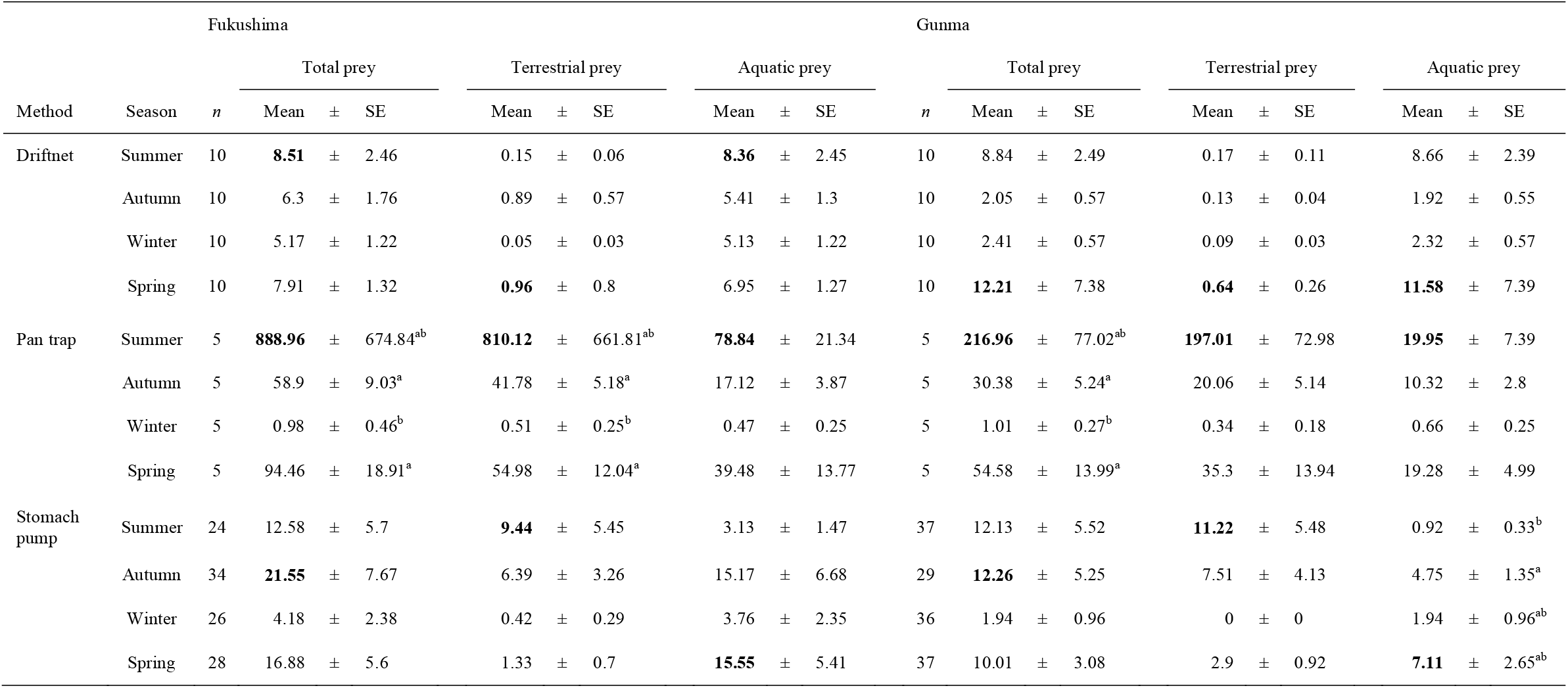
Availability of total, terrestrial, and aquatic prey for charr estimated using driftnets (mg/m^3^), pan traps (mg/m^2^/day), and stomach pumps (mg/individual) at each site in each season. Data are presented as the mean biomass ± standard error (SE). Bold characters denote the maximum mean biomass of each prey type (total, terrestrial, and aquatic) in each season. Different alphabetical letters indicate statistically significant differences by post hoc comparisons (*P* < 0.05).

**TABLE 3.** Results of permutation analyses of variance testing for effect of season on biomass of total, terrestrial, and aquatic preys estimated by driftnet, pan trap, and stomach pump in each site. Bold characters indicate statistically significant differences (*P* < 0.05).

| Method | Prey type | Fukushima |  | Gunma |  |
| --- | --- | --- | --- | --- | --- |
|  |  | *F* | *P* | *F* | *P* |
| Driftnet | Total prey | 0.75 | 0.54 | 1.62 | 0.12 |
|  | Terrestrial prey | 0.96 | 0.42 | 3.17 | **0.01** |
|  | Aquatic prey | 0.83 | 0.46 | 1.50 | 0.15 |
| Pan trap | Total prey | 1.55 | **<0.01** | 7.88 | **<0.01** |
|  | Terrestrial prey | 1.39 | **<0.01** | 7.58 | **<0.01** |
|  | Aquatic prey | 6.97 | **<0.01** | 4.48 | **0.02** |
| Stomach<br>pump | Total prey | 1.52 | 0.22 | 1.47 | 0.22 |
|  | Terrestrial prey | 1.78 | 0.15 | 2.18 | 0.07 |
|  | Aquatic prey | 1.83 | 0.14 | 3.18 | **<0.01** |

**TABLE 4.** Total biomass (mg) of terrestrial and aquatic prey taxa estimated from the stomach contents of charr.

| Fukushima |  |  |  | Gunma |  |  |  |
| --- | --- | --- | --- | --- | --- | --- | --- |
| Terrestrial |  | Aquatic |  | Terrestrial |  | Aquatic |  |
| Anura | 178.70 | Isopoda | 380.44 | Orthoptera | 297.63 | Trichoptera | 134.93 |
| Orthoptera | 111.24 | Decapoda | 318.09 | Coleoptera | 190.25 | Diptera | 125.08 |
| Oligochaeta | 75.40 | Plecoptera | 117.10 | Anura | 120.70 | Decapoda | 74.30 |
| Diplopoda | 35.00 | Diptera | 112.77 | Lepidoptera | 43.20 | Ephemeroptera | 70.72 |
| Coleoptera | 21.50 | Trichoptera | 87.91 | Diplopoda | 32.90 | Salmoniformes | 29.10 |
| Unknown | 20.77 | Odonata | 44.30 | Araneae | 26.53 | Odonata | 26.32 |
| Hemiptera | 18.92 | Ephemeroptera | 43.58 | Hymenoptera | 11.83 | Plecoptera | 24.55 |
| Hymenoptera | 15.56 | Coleoptera | 19.58 | Oligochaeta | 7.20 | Coleoptera | 10.88 |
| Araneae | 8.53 | Neuroptera | 0.01 | Unknown | 5.41 | Isopoda | 8.70 |
| Gastropoda | 5.90 |  |  | Hemiptera | 4.30 | Neuroptera | 0.17 |
| Collembola | 0.54 |  |  | Collembola | 0.23 |  |  |
| Acari | 0.03 |  |  | Acari | 0.05 |  |  |

### 3.2 ^137^Cs concentrations in litter and prey

The ^137^Cs concentrations in terrestrial and aquatic litter from the Fukushima site were 14,700–36,000 and 3,960–8,500 Bq/kg, respectively, whereas those from the Gunma site were 2,620–8,830 and 1,100–7,320 Bq/kg, respectively (Table S1). ^137^Cs concentrations in terrestrial litter were significantly higher than those in aquatic litter at both sites, and the difference in ^137^Cs concentrations between aquatic and terrestrial litter was greater at the Fukushima site than at the Gunma site (Figure 2a,b).

**FIGURE 2.**
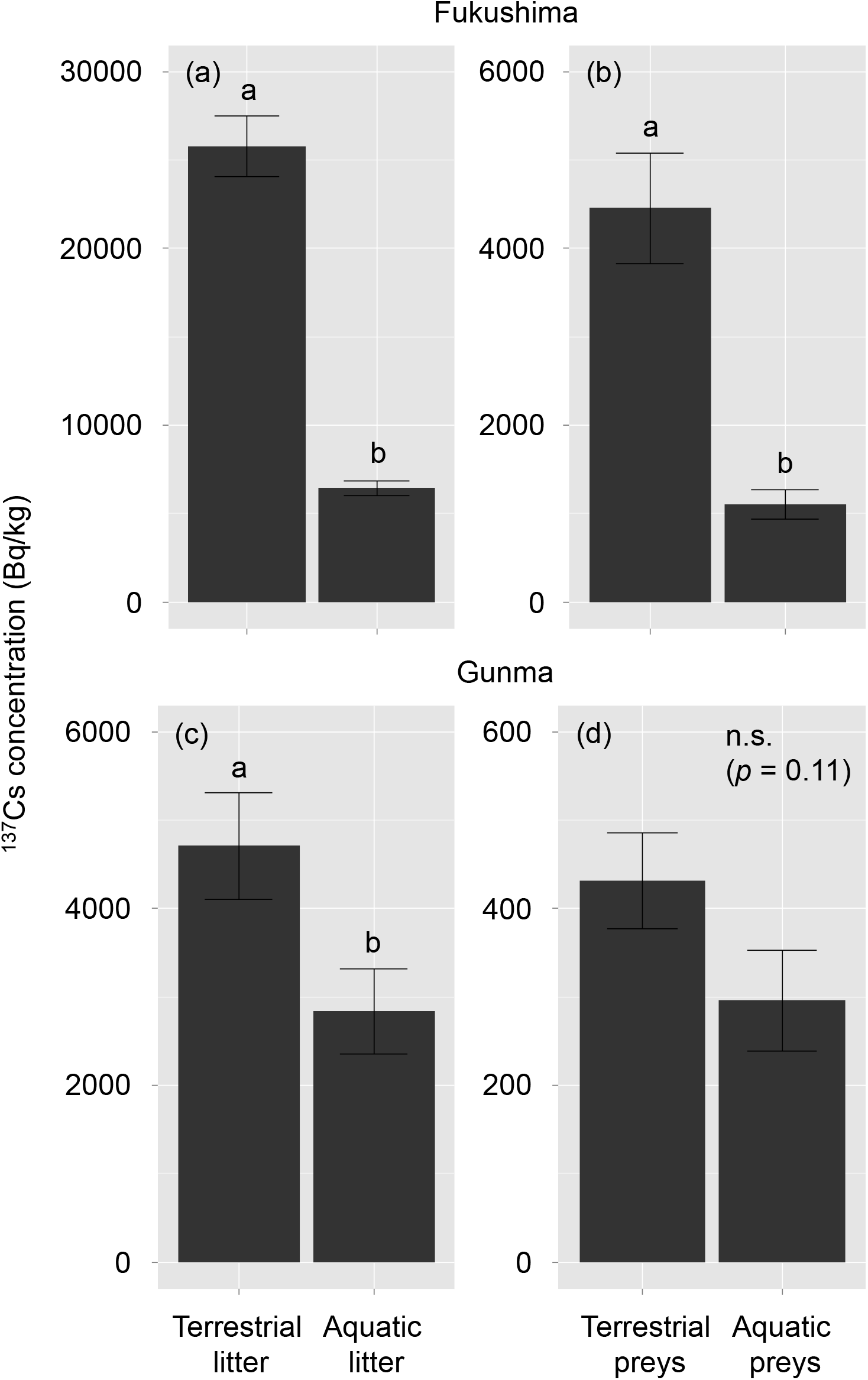
Mean ^137^Cs concentrations in Japanese cedar litter and preys collected from both terrestrial and aquatic habitats in the Fukushima (a and b) and Gunma (c and d) sites. Error bars indicate ± standard error. The data obtained from all the sampling periods was pooled in this figure and each value can be referenced from Table S1. Different alphabetical letters indicate statistically significant difference (permutation analyses of variance, *P* < 0.05) and n.s. indicates insignificant difference.

We measured ^137^Cs concentrations in four orders of terrestrial organisms (*n* = 22) and eight orders of aquatic organisms (*n* = 31) from the Fukushima site, and four orders of terrestrial organisms (*n* = 16) and six orders of aquatic organisms (*n* = 22) from the Gunma site (Table S1). For both sites, these measurements included the three most common taxa found in the stomach contents of charr, except terrestrial Oligochaeta at the Fukushima site. In total, the Cs measurements of terrestrial and aquatic prey included 88.1% of the stomach contents of charr at the Fukushima site and 87.6% of those at the Gunma site (Table 4).

The ^137^Cs concentrations in prey were significantly higher in terrestrial than aquatic environments at the Fukushima site but not at the Gunma site (Figure 2c,d). The median ^137^Cs concentrations in terrestrial and aquatic prey were 4,000 and 853 Bq/kg, respectively, at the Fukushima site and 393 and 274 Bq/kg at the Gunma site. The estimated ^137^Cs concentrations in prey at the Fukushima site changed seasonally (Table 5 and Figure 3a) by seasonal shifts of the relative contribution of terrestrial and aquatic prey (Table 2) whereas those remained relatively constant across seasons at the Gunma site due to the greater similarity in ^137^Cs concentrations in terrestrial and aquatic prey (Figure 3b).

**TABLE 5.** Results of permutation analyses of variance testing for effect of season on estimated ^137^Cs concentrations in prey, specific metabolic rate and ^137^Cs concentrations in charr. Bold characters indicate statistically significant differences (*P* < 0.05).

|  | Fukushima |  | Gunma |  |
| --- | --- | --- | --- | --- |
|  | *F* | *P* | *F* | *P* |
| Estimated $^{137}\text{Cs}$ concentrations in prey | 5.32 | **<0.01** | 6.41 | **<0.01** |
| Specific metabolic rate | 128.89 | **<0.01** | 151.63 | **<0.01** |
| $^{137}\text{Cs}$ concentrations in charr | 0.02 | 1.00 | 4.35 | **0.02** |

**FIGURE 3.**
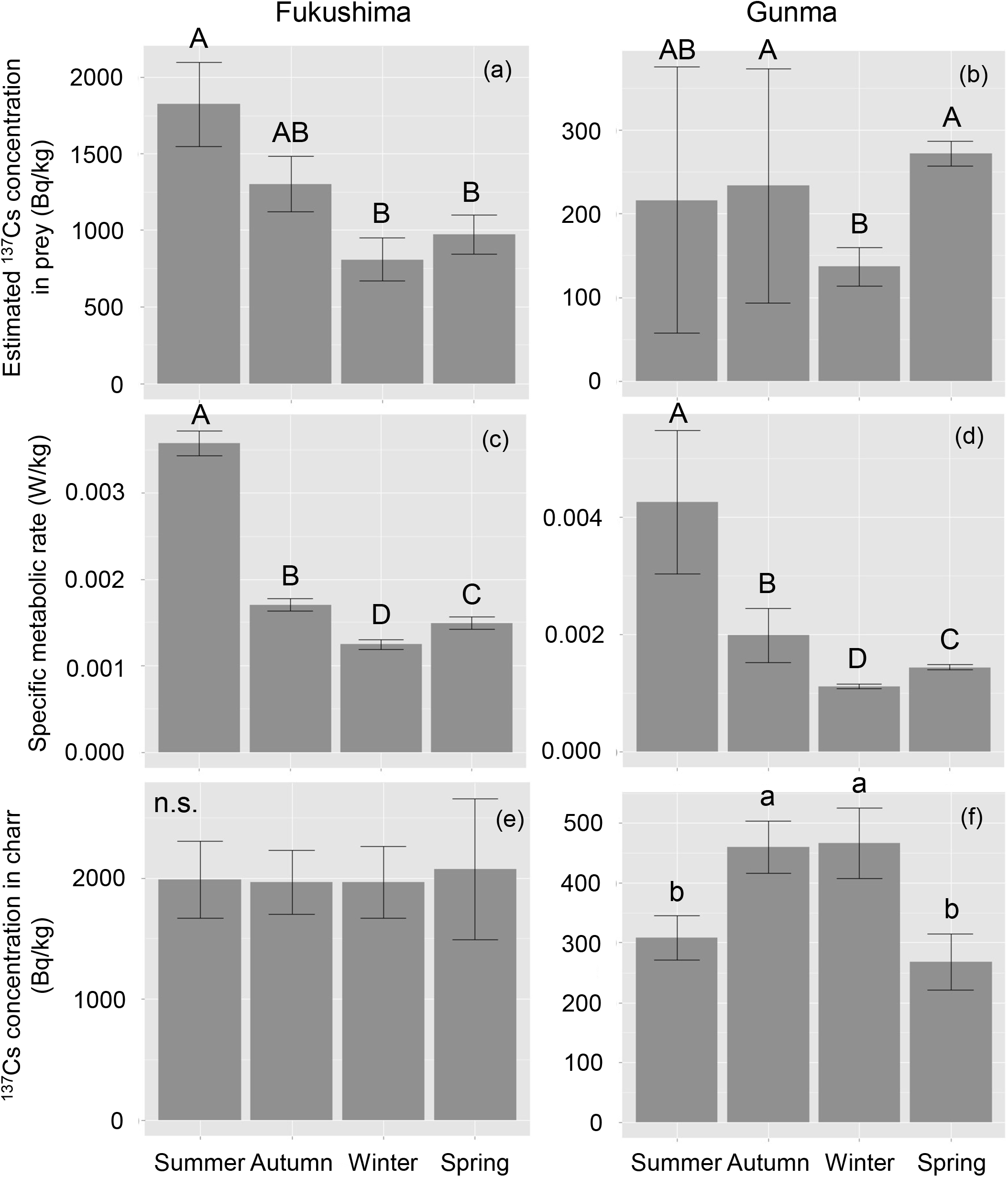
Seasonal variations in the estimated ^137^Cs concentrations in prey, specific metabolic rates of charr and ^137^Cs concentrations in charr at the Fukushima (a, c, and e) and Gunma sites (b, d, and f). Error bars indicate ± standard error. Different alphabetical letters indicate statistically significant difference (post hoc comparisons, *P* < 0.05 for uppercase and < 0.1 for lowercase) and n.s. indicates insignificant difference. Summer: August, autumn: November, winter: February, spring: May.

### 3.3 Charr metabolism and ^137^Cs concentration

Because of the structure of the equation of specific metabolic rate, the specific metabolic rates of charr increased with stream water temperature and were thus highest in summer and lowest in winter at both sites (Table 5 and Figure 3c,d). The ranges of the specific metabolic rates of charr were: 0.0008–0.0041 and 0.0006–0.0070 W/kg at the Fukushima and Gunma sites, respectively.

The concentrations of ^137^Cs in charr ranged from 1,190 to 6,080 and from 118 to 617 Bq/kg at the Fukushima and Gunma sites, respectively (Table S1). The ANOVA models showed that the ^137^Cs concentrations in charr did not differ among seasons at the Fukushima sites (Table 5 and Figure 3e) whereas those at the Gunma site was relatively higher in autumn and winter (Table 5 and Figure 3f).

## 4 DISCUSSION

The charr at our study sites consumed both terrestrial and aquatic organisms, and the consumption amounts of terrestrial and aquatic preys were greater in summer and spring, respectively. This pattern was in agreement with seasonal dependencies in terrestrial and aquatic food resources for salmonids in other temperate forested streams (Baxter et al., 2005; Nakano et al., 1999a). The seasonal dependencies were basically reported from broad-leaved deciduous forests (e.g., Cloe & Garman, 1996; Kawaguchi & Nakano, 2001; Nakano & Murakami, 2001), but our results confirmed that the similar seasonal patterns can occur also in evergreen conifer plantations.

The ^137^Cs concentrations in Japanese cedar litter were significantly higher in terrestrial than in aquatic environments as previously reported (Gomi et al., 2018; Sakai et al., 2015). The concentration difference in both litter and preys was greater at the Fukushima site than at the Gunma site. The results suggest that ^137^Cs leaching from litter can extensively induce ^137^Cs concentration difference in preys in between terrestrial and aquatic environments (Sakai et al., 2016a). Importantly, the ^137^Cs concentration difference in between terrestrial and aquatic preys becomes greater in more contaminated sites (cf. Sakai et al., 2016b).

Because charr consumed terrestrial prey more in summer than in other seasons, the estimated ^137^Cs concentration in prey was peaked in summer at the Fukushima site. Meanwhile, such peak was not observed at the Gunma site, as ^137^Cs concentration was relatively similar between terrestrial and aquatic preys. Therefore, the highest availability of terrestrial prey in summer can cause more ^137^Cs uptake to charr in more contaminated headwaters. To validate the pattern observed here, grasping the seasonality of ^137^Cs concentration in prey for headwater salmonids at various ^137^Cs deposition levels is required for future studies.

In contrast, seasonal change in metabolism of charr was commonly high in summer and low in winter at the sites because that is basically controlled by water temperature (Brown, 2004). At the Fukushima site, because both ^137^Cs concentrations in prey and specific metabolic rates of charr peaked in summer, ^137^Cs uptake and elimination in charr could have cancelled out any seasonal ^137^Cs fluctuations, and thus ^137^Cs concentrations in charr were stable. On the other hand, fluctuation in specific metabolic rate was likely related to the ^137^Cs concentrations in charr at the Gunma site because ^137^Cs concentration in preys was consistent. Therefore, the ^137^Cs concentrations in charr could be higher in winter. These results suggest a possible threshold at which uptake or elimination influences ^137^Cs concentrations in charr depending on initial ^137^Cs fallout volumes.

The result also indicated that the ^137^Cs concentrations in charr were higher in autumn at the Gunma site. This may be related to ^137^Cs uptake through autumn inputs of Rhaphidophorid insects, which are of most contaminated invertebrates in forests (Sakai et al., 2016a), into streams. While Rhaphidophorids were actually the most abundant contents of charr guts at the Gunma site, dive of the insects from riparian forests to streams generally peaks from September to October (Sato et al., 2011, 2012). Thus, our charr samples in autumn that collected in November might have already consumed substantial volumes of the highly contaminated subsidy. Because effective half-life of ^137^Cs was approximately two months for *S. leucomaenis* (Matsuda et al., 2020), impact of the resource subsidy on ^137^Cs accumulation in charr could prolong to late autumn. The relationship between Rhaphidophorid inputs and ^137^Cs contamination in headwater salmonids would be an emerging topic that critically related to ^137^Cs dynamics in headwater stream ecosystems.

Although this study could not estimate quantitative ^137^Cs input/output in charr, biomass of prey consumed by charr can influence ^137^Cs concentrations in the fish (Forseth et al., 1991; Ugedal et al., 1997). For example, charr in the Gunma site consumed more prey in autumn than in spring suggests that ^137^Cs uptake was higher in autumn. This may also be an important process that caused higher ^137^Cs concentrations in charr in autumn than in spring irrespective of the higher specific metabolic rate in autumn. Meanwhile in the Fukushima site, the ^137^Cs uptake and elimination rates might be balanced enough to induce the seasonal consistency of ^137^Cs concentrations in charr. This raises an assumption that charr inhabited in more contaminated sites than in the Fukushima site may show peaks of ^137^Cs concentration in summer due to excessive ^137^Cs uptake through contaminated terrestrial prey consumptions. Developing a method estimating quantitative ^137^Cs input/output in fishes is required for future studies.

This study was conducted in an early stage of the radiation contamination (one to two years after the accident). Currently, most litter is less contaminated compared to those in the early stage because leaves and needles contaminated by the ^137^Cs fallout in 2011 had already been abscised. In particular, ^137^Cs concentrations in Japanese cedar litter was remarkably decreased with time, whereas those in deciduous broad-leaved litter remained at higher levels through ^137^Cs root uptake (Onda et al., 2020). Therefore, difference of ^137^Cs concentrations in litter between forest and stream ecosystems may become smaller especially in coniferous plantations of heavily contaminated areas, and current seasonal fluctuations in ^137^Cs concentration in headwater salmonids may be influenced primarily by their metabolism. These characteristics of temporal changes in ^137^Cs concentration in litter should be crucial when assess ^137^Cs uptake to fishes from preys that exhibit variable ^137^Cs concentrations through the leaching effects on detrital food webs between forest and stream ecosystems.

In this study, both metabolism and ^137^Cs concentrations in prey items, whose seasonality are varied due to initial ^137^Cs fallout volume, were expected to be important determinants for ^137^Cs concentrations in charr. Meanwhile, the difficulty on radioactivity measurements for small-volume samples has largely hindered quantitative (not concentration) assessments on ^137^Cs flows in predator-prey interactions. Our results also could not estimate quantitative input and output of ^137^Cs in fishes, but the plausible processes proposed here suggest the importance of the ^137^Cs input/output. At least, our results suggested that less ^137^Cs-concentrated freshwater salmonids might be more obtainable in hotter season, when specific metabolic rate is higher, in moderately contaminated sites. Meanwhile in highly contaminated sites, reducing terrestrial preys belonging to heavily contaminated detrital food webs is important to acquire less contaminated fishes. Further studies on radiocesium transfers and dynamics in fishes would help develop better contamination managements in freshwater fisheries. Then, temporal changes in ^137^Cs concentrations in litter is crucial to assess headwater salmonids that belong to detrital food webs tangled in forest and stream ecosystems (Sakai et al., 2021).

## ACKNOWLEDGMENTS

We thank the members of the Watershed Hydrology and Ecosystem Management Laboratory at the Tokyo University of Agriculture and Technology for their support in conducting the fieldwork. In addition, we extend our appreciation to the members of the Watershed Conservation and Management Laboratory at Hokkaido University. Insightful comment for the improvement of this manuscript was provided by anonymous editors and reviewers. This study was supported by the Environmental Research Fund (ZD-1202) of the Ministry of the Environment, Japan, and the JSPS KAKENHI Grant Numbers 24248058 and 15H04511. The authors declare that they have no conflicts of interest.

